# Sexual selection and inbreeding: two efficient ways to limit the accumulation of deleterious mutations

**DOI:** 10.1101/273367

**Authors:** E. Noël, E. Fruitet, D. Lelaurin, N. Bonel, A. Ségard, V. Sarda, P. Jarne, P. David

## Abstract

Theory and empirical data showed that two processes can boost selection against deleterious mutations, thus facilitating the purging of the mutation load: inbreeding, by exposing recessive deleterious alleles to selection in homozygous form, and sexual selection, by enhancing the relative reproductive success of males with small mutation loads. These processes tend to be mutually exclusive because sexual selection is reduced under mating systems that promote inbreeding, such as self-fertilization in hermaphrodites. We estimated the relative efficiency of inbreeding and sexual selection at purging the genetic load, using 50 generations of experimental evolution, in a hermaphroditic snail (*Physa acuta*). To this end, we generated lines that were exposed to various intensities of inbreeding, sexual selection (on the male function) and nonsexual selection (on the female function). We measured how these regimes affected the mutation load, quantified through the survival of outcrossed and selfed juveniles. We found that juvenile survival strongly decreased in outbred lines with reduced male selection, but not when female selection was relaxed, showing that male-specific sexual selection does purge deleterious mutations. However, in lines exposed to inbreeding, where sexual selection was also relaxed, survival did not decrease, and even increased for self-fertilized juveniles, showing that purging through inbreeding can compensate for the absence of sexual selection. Our results point to the further question of whether a mixed strategy combining the advantages of both mechanisms of genetic purging could be evolutionary stable.

## Background

Natural selection is often considered as an improvement process, progressively increasing the adaptation of organisms to their environment. However, what is known about the distribution of the fitness effects of mutations indicates that they are rarely advantageous, and mostly vary from slightly deleterious to lethal [1,2]. Thus, natural selection is more often purifying genomes from deleterious mutations than accumulating adaptive changes. Characterizing when and why this genetic purging is efficient, is therefore crucial to understanding the evolutionary fate of both wild and captive populations [3].

The efficiency of genetic purging depends on mutations being expressed at the phenotypic level on which selection is acting. Two processes are considered to enhance this expression, acting in different ways. The first is sexual selection [4,5]. Indeed, if males with large numbers of mutations have low reproductive success (e.g., because of female mate choice or male-male competition), mutations are eliminated from the population with little demographic cost. This idea relies on the genic capture hypothesis [6], *i.e*. that sexually selected traits are highly condition-dependent, so that many deleterious alleles will tend to pleiotropically affect male success. A large number of experiments have been conducted to test the effect of sexual selection on purging ([7–13] and previous ones reviewed in [14]). For example, Lumley et al. (2015) recently showed that the suppression of sexual selection for many generations through imposed monogamy or highly female-biased sex-ratios resulted in elevated genetic loads in flour beetles. However, this type of studies yielded overall inconsistent results [14]. This inconsistency may be due to sexual-conflict alleles present in the initial standing variation of populations. Indeed monogamy relaxes not only sexual selection but also sexual conflict, yielding benefits that may conceal the disadvantages of suppressing sexual selection [14–18]. This can be avoided by tracking the fate of a limited number of purely deleterious mutations introduced, induced or accumulated *de novo* in populations [7,13,17,19–22] instead of the standing load. These mutations, however, represent a small fraction of the genome (and therefore a small expected cumulative effect) compared to the typical standing mutation load of natural populations.

The second process that can purge the genetic load is inbreeding [23,24], especially under its strongest form (*i.e*. self-fertilisation). Inbreeding increases homozygosity, exposing recessive deleterious alleles to natural selection. It has been argued, however, that only part of the load, that is, highly recessive alleles with large effect, can be purged quickly [23,25]. Inbred populations still tend to accumulate slightly deleterious, partially-recessive mutations, a process aggravated by selective interference and the decrease of effective population size associated with inbreeding [26].

Self-fertile hermaphroditic organisms can be exposed to both strong sexual selection and self-fertilization, depending on how they reproduce, and can purge their genetic load through both processes. Many hermaphroditic animals or plants are actually polyandrous and subject to intense sexual selection [27–32], and some of them also have the ability to self-fertilize. Under reduced mate or pollen availability, these populations may self-fertilize as a reproductive assurance strategy [33], and some lineages eventually evolve towards preferential, often nearly-complete, self-fertilization [34,35]. Evolutionary transitions towards self-fertilization entail a relaxation of sexual selection that may compromise the purging of deleterious mutations. To understand how the mutation load affects the evolution and persistence of selfing lineages, it is important to evaluate the relative efficiency of inbreeding and sexual selection at promoting genetic purging. A related issue is whether the two processes affect the same pool of deleterious mutations. As mentioned above, purging by inbreeding depends on the homozygous effect (and recessivity) of mutations on survival and fertility, while sexual selection predominantly affects mutations with large heterozygous effects on male competitiveness. If these two categories do not completely overlap, both the quantity and the nature of the purged mutations may differ, those purged by inbreeding being on average more recessive. It is therefore important to monitor how these two processes affect not only fitness traits, but also inbreeding depression, which reflects the recessive load. Such studies have not been conducted to the best of our knowledge.

We conducted such a comparison in *Physa acuta*, a self-compatible hermaphroditic freshwater snail. This species shows both pre- and post-copulatory sexual selection on the male function [30,36,37] but not -or very little- on the female function. This is because male reproductive success is significantly increased by remating after the first copulation, while female reproductive success is not. We generated experimental evolution lines of *P. acuta* belonging to four types. The first three types were purely outcrossing; they included the control type (maintained under a polygamous mating system typical of high-density populations) and two types with relaxed selection on respectively the female or male function (selection on the male function being relaxed by removing sexual selection). The fourth type was undergoing regular self-fertilization and no sexual selection. We compared the accumulation of deleterious mutations under these four regimes by following the change in juvenile survival in selfed and outcrossed offspring from all lines in common garden experiments at two evolutionary time points (20 and 50 generations). Juvenile survival was chosen because it is an important component of fitness and is not sex-specific.

We expected, if sexual selection enhances purging, that juvenile survival would decrease in lines with relaxed sexual selection, and that this decrease would be stronger than that observed in lines with relaxed selection on female reproduction. We expected, if inbreeding also promotes purging, that juvenile survival would be higher in lines experiencing both frequent self-fertilization and relaxed sexual selection, than in those with only relaxed sexual selection. In addition, we expected purging by inbreeding to target more specifically deleterious alleles with recessive effect, resulting in a reduced inbreeding depression in lines exposed to inbreeding compared to all others.

## Methods

### Experimental evolution

Our model species, *Physa acuta* (Hygrophila, Physidae), is a simultaneous hermaphroditic snail that naturally outcrosses but can self-fertilize its eggs in the absence of mates [38]. Experimental evolution lines (hereafter EELs) were founded in 2008, from a common, genetically diverse pool of individuals issued from 10 natural populations sampled in November 2007 near Montpellier, France. Details on the common source population, as well as on usual rearing procedures for this species, can be found in [39].

We created four types of EELs (C, M, F, and S) allowing us to impose different intensities of sexual selection (on the male function), different intensities of selection on female reproduction, and different levels of inbreeding. To this end, we manipulated both the mating system and the among-individual variance in female reproductive success (Table 1; supplementary Fig. S1). Three mating systems were used: (i) mating in large groups maintained for one week (hereafter mass-mating), which in this species ensures that each individual is inseminated by several partners and that competition occurs among different male-acting individuals to obtain mates, and among stored sperm from different donors to fertilize eggs (Pélissié et al. 2012, 2014). Thus, this system ensures 100% outcrossing, and authorizes sexual selection. (ii) mating in random monogamous pairs for one week, which relaxes sexual selection, but still under outcrossing (iii) self-fertilization, which relaxes sexual selection and imposes inbreeding. Variance in female reproductive success was manipulated when collecting juveniles laid separately by all mothers in individual boxes (supplementary Figure S1). In the “no regulation” treatment, we pooled together all juveniles to constitute a pool from which the next generation of adults was drawn at random, thus incorporating the natural variance in female reproductive success, and letting selection on female reproduction operate. In the “juvenile-regulation” treatment we collected a fixed number of juveniles per mother, irrespective of how many were present, therefore limiting the variance in female reproductive success and the selection on female reproduction. Note that this selection is fertility selection and not of sexual nature; it has been previously shown that sexual selection (as measured by the Bateman gradient) acts only on the male function in this species (Pélissié et al. 2012).

**Table 1:** Selective pressures in the different types of experimental evolutionary lines.

| Line type | Mating system | (Sexual) selection on the male function | (Nonsexual) selection on the female function | Inbreeding |
| --- | --- | --- | --- | --- |
| Control (C) | Mass-mating | Yes | Yes | No |
| Male-selected (M) | Mass-mating | Yes | No | No |
| Female-selected (F) | Monogamy | No | Yes | No |
| Selfing (S) | Selfing and monogamy | No | Yes | Yes |

These treatments were combined as follows to produce the four types of lines (Table 1; supplementary Fig. S1). In the control type (C) both male and female selection was operating since individuals reproduced in mass mating and juvenile numbers were not regulated. In the male-selected type (M), snails reproduced in mass-mating, inducing sexual selection on the male function, but female selection was relaxed through juvenile regulation. The opposite occurred in the female-selected type (F): male sexual selection was relaxed by imposing monogamy, but juveniles were not regulated, preserving selection on the female function. The situation in the selfing type (S) was a bit more complex: the mating system alternated between selfing (odd-numbered generations) and monogamy (even-numbered generations) and juvenile numbers were never regulated. The selection regime in S lines therefore resembled that of F lines (no sexual selection, but selection on female reproduction is preserved), except that the s-lines were regularly exposed to inbreeding. We did not enforce selfing at each generation in S-lines, based on previous work suggesting that too many lines would have been lost given the strong inbreeding depression [40]. The practical details of the passage of generations in each line type are given in supplementary Fig. S1. Each line type was replicated twice (e.g., C1 and C2 for the C type), resulting in eight EELs in total, all evolving at a population size of ca. 80-90 reproductive adults per generation.

### Juvenile survival assay

The EELs were initiated in 2008, and we sampled them twice, around the 20th (year 2012) and 50th (2016) generations of experimental evolution, to evaluate change in juvenile survival. We chose juvenile survival as our focal fitness component because it affects male and female fitness in the same way in simultaneous hermaphrodites, and therefore reflects the load of non sex-specific deleterious mutations. In addition, we evaluated the survival of both outcrossed and selfed offspring in order to compare the magnitude of inbreeding depression generated by deleterious mutations among lines, which depends on their degree of recessivity.

Individuals used as parents in the first experiment (2012) were extracted from generations G17 (S1 and S2), G22 (M2 and F2) and G23 (other lines). The second experiment took place in 2016-2017 and was spread over a year in order to include a large number of individuals. We therefore extracted individuals from each EEL at different successive generations (C1: G47-G52, C2: G48-G51, M1: G51-G52, M2: G50-G51, F1: G51-G53, F2: G50-G51, S1:G42-G47, S2: G43-47). However, we assume that evolution of survival is slow enough (Noel et al. 2016) to consider that this experiment constitutes a single evolutionary time point, near the 50^th^ generation. In both experiments, the generations studied differ among lines, because the EELs slowly got desynchronized since their foundation. This is especially true in S lines because isolated mature adults wait for ca. two weeks before selfing compared to the moment they would engage in outcrossing [41], resulting in a longer generation time in S lines than in others. Furthermore, we always started the experiments with outcrossed parents. Therefore, assays were possible only in even-numbered generations for the S lines (extending the total time span of experiments). Each experiment was subdivided in several temporal blocks (based on the date of egg collection) to account for the possible effect of temporal variation in raising conditions on juvenile survival (4 and 13 blocks respectively for the two experiments). Note that some data (C and S lines from the first experiment) have already been used in a previous study[39]. In what follows, we refer to the first and second experiment as the G20 and G50 experiment respectively.

For each experiment, Half of the individuals extracted from the lines were outcrossed either in pairs (G20 experiment) or in mass-mating (G50) during three days, and then re-isolated and allowed to lay eggs for three days. The other half remained isolated until they started to lay self-fertilized eggs, and then allowed to lay for three more days. We collected the clutches in both cases, counted the eggs, and added food. Fifteen days later, the juveniles alive were counted in each box. Juvenile survival was computed as the number of juveniles divided by the initial number of eggs. Eggs hatch in approximately one week, so the eggs that failed to produce live juveniles either did not hatch, or produced hatchlings that died during their first week. In total, we obtained clutches from 3328 different individuals (see Table 3 for their repartition among experiments, treatments and line types), resulting in 234,187 eggs and 104,540 live juveniles.

**Table 3.** Estimates, effect sizes and tests of model terms in the complete linear model (same GLMM as in Table 2). Abbreviations for effects: L = Line type, T= treatment, E = Experiment. Sample size (number of individuals laying eggs), predicted and raw means of juvenile survival are provided by group, each group being characterized by a combination of line type (C, M, F, or S), treatment (outcrossed or selfed) and experiment (G20 or G50). By construction, the mean of the reference group (C/outcrossed/G20) is predicted by the intercept of the model. the factor levels (or combinations thereof) being tested are highlighted. Model predictions (de-logited) differ from raw means because of the use of logit scale in the model, and the presence of random factors to model means and variance within groups (the logit scale gives more weight to values close to 0 or 1). We therefore provide both de-logited model estimates and raw means. Effect size (model term divided by its SE) and associated Wald-tests are given together with their interpretation. *P*-values have been corrected for multiple-testing by the Benjamini-Hochberg FDR procedure, when three line types (M, F, S) were separately compared with controls (Line type effects and their interactions). *P* values <0.05 are in bold. ID=inbreeding depression.

| Model term | Group of individuals concerned | N | Survival (predicted) | Survival (raw means ± SE) | Effect size | *P* | Interpretation |
| --- | --- | --- | --- | --- | --- | --- | --- |
| Intercept | C/outcrossed/G20 | 137 | 0.492 | 0.545± 0.022 | | | |
| T (selfed) | C/**selfed**/G20 | 77 | 0.201 | 0.293± 0.027 | -5.680 | **1×10<sup>-8</sup>** | ID is significant in C lines in G20 |
| E (G50) | C/outcrossed/**G50** | 445 | 0.557 | 0.602 ±0.013 | 0.732 | 0.464 | Survival of C lines under outcrossing does not change between G20 and G50 |
| T*E(selfed, G50) | C/**selfed/G50** | 404 | 0.257 | 0.319± 0.014 | 0.177 | 0.860 | ID in C lines does not change between G20 and G50 |
| L (M) | **M**/outcrossed/G20 | 45 | 0.322 | 0.374±0.049 | -2.506 | **0.018** | M lines survive less than C lines under outcrossing in G20 |
| L (F) | **F**/outcrossed/G20 | 44 | 0.237 | 0.293±0.034 | -3.972 | **2×10<sup>-4</sup>** | F lines survive less than C lines under outcrossing in G20 |
| L (S) | **S**/outcrossed/G20 | 133 | 0.438 | 0.496±0.019 | -1.157 | 0.247 | Survival does not differ between S and C lines under outcrossing in G20 |
| L*T(M,selfed) | **M/selfed**/G20 | 38 | 0.156 | 0.226±0.035 | 0.936 | 0.523 | ID does not differ between C and M lines in G20 |
| L*T(F,selfed) | **F/selfed**/G20 | 34 | 0.090 | 0.151± 0.032 | 0.462 | 0.644 | ID does not differ between C and F in G20 |
| L*T(S,selfed) | **S/selfed**/G20 | 84 | 0.406 | 0.459±0.028 | 3.738 | **6×10<sup>-4</sup>** | ID is lower in S than in C lines in G20 |
| L*E(M,G50) | **M**/outcrossed/**G50** | 268 | 0.619 | 0.619±0.015 | 2.477 | **0.020** | Survival of outcrossed juveniles differs between C and M lines in G20, but not in G50 |
| L*E(F,G50) | **F**/outcrossed/**G50** | 234 | 0.102 | 0.191±0.014 | -3.200 | **0.004** | The difference in survival between F and C lines increases between G20 and G50, under outcrossing |
| L*E(S,G50) | **S**/outcrossed/**G50** | 456 | 0.541 | 0.569±0.012 | 0.480 | 0.631 | Survival of outcrossed juveniles is stable in time for both S and C lines |
| L*T*E(M,selfed,G50) | **M/selfed/G50** | 198 | 0.304 | 0.372±0.018 | -0.800 | 0.424 | ID is similar between M and C in both G50 and G20 |
| L*T*E(F,selfed,G50) | **F/selfed/G50** | 261 | 0.070 | 0.161± 0.012 | 1.251 | 0.316 | ID is similar between F and C in both G50 and G20 |
| L*T*E(S,selfed,G50) | **S/selfed/G50** | 469 | 0.355 | 0.408±0.012 | -1.579 | 0.316 | The difference in ID between C and S lines is preserved (or reduced but not significantly) between G20 and G50 |

**Table 2.** Results of the linear model (GLMM) on juvenile survival (Binomial variable with logit link, with individual observation as a random factor to account for overdispersion). Tests of fixed effects by Likelihood-ratio chi-square tests (LRT) are provided; the number of degrees of freedom is 1 except for line type and interactions that involve line type (3 df). *P*-values <0.05 are in bold. To facilitate interpretation, we laid out the question and answers associated to each test of fixed effect, and the percentage of total variance (rather than raw values) explained by each random effect. ID=inbreeding depression.

| Question | Effect | LRT<br>($\chi^2$) | *P* | Interpretation of tests for fixed effects (% explained variance for random effects) |
| --- | --- | --- | --- | --- |
| **FIXED EFFECTS** |  |  |  |  |
| Is there a difference in average survival between outcrossed and selfed juveniles, reflecting ID? | Treatment | 21.74 | **$3 \times 10^{-6}$** | Yes, ID is significant overall |
| Is there a change in average juvenile survival over generations, between G20 and G50? | Experiment | 0.203 | 0.652 | No, survival does not significantly change between G20 and G50 |
| Is there a change in average impact of inbreeding (ID) over generations? | Treatment * experiment | 0.135 | 0.714 | No, average ID does not significantly change between G20 and G50 |
| Does average juvenile survival differ depending on selection regime? | Line type | 25.25 | **$1.4 \times 10^{-5}$** | Yes, line types significantly differ in their average survival rate |
| Does ID differ depending on selection regime? | Line type * treatment | 7.821 | **0.050** | Yes, ID significantly differs among line types |
| Does the selection regime affect the temporal change in survival between G20 and G50? | Line type * experiment | 15.215 | **0.002** | Yes, survival changes between G20 and G50 in a line type-specific way |
| Does the selection regime affect the temporal change in ID between G20 and G50? | Line type * treatment * experiment | 5.501 | 0.139 | No, there are no significant line-specific changes in ID between G20 and G50 |
| **RANDOM EFFECTS** |  |  |  |  |
| | Block | 89.789 | $< 2 \times 10^{-16}$ | 3.02% |
|  | Replicate (within line type) | 2.256 | 0.133 | 0.11% |
|  | Replicate*treatment | 10.956 | **0.012** | 0.40% |
|  | Replicate*experiment | 1.812 | 0.612 | 0.01% |
|  | Replicate*treatment *experiment | 1.811 | 0.770 | 0.08% |
|  | Individual observation (residual) |  |  | 66.33% |

### Statistical analyses

Juvenile survival (coded as a two-column vector, number of juveniles alive and number of eggs that failed to produce juveniles) was analysed as a binomial variable using a Generalized Linear Mixed Model with the lme4 package in R [42]. Experiment (G20 vs G50), line type (C, M, F, S) and treatment (outcrossed vs selfed), and their interactions, were included as fixed effects. Replicate within line type, its interactions with fixed effects (other than line type), and temporal block, were added as random effects. Moreover, we added individual identity as a random factor in order to correct for overdispersion [43]. The fixed and random effects were tested using chi-square likelihood-ratio tests (LRT). In addition we used Wald tests, for each fixed factor, to test which levels of fixed factors were significantly different from the reference level (controls), correcting for multiple testing by the FDR method of Benjamini-Hochberg over the different levels [44].

### Microsatellite data

We measured genetic diversity at molecular markers to assess whether line types underwent different amounts of neutral genetic drift, and whether the individuals produced by crosses were indeed cross-fertilised as we intended (i.e. that none of the lines, including the S type, evolved some form of preferential self-fertilization). We assessed variation at seven microsatellite loci (AF762, AF764, Pac1, Pac2, Pasu11, Pasu2, Pasu9; protocols in [45,46]) in 32 individuals from the 49^th^ generation of each EEL. Note that for the S lines, the individuals we genotyped were actually offspring from crosses between the G48 parents (instead of the selfed offspring that were used to propagate the lines). Microsatellite variation was quantified using Nei’s unbiased estimate of genetic diversity (*H*_*e*_), observed heterozygosity (*H*_*o*_), the number of alleles per locus, and Weir & Cockerham’s (1983) estimator (*f*) of the inbreeding coefficient F_IS_, using the software Genetix [47].

## Results

Depending on treatment, experiment and line type, survival varied between 0.201 (F, G20, selfed) and 0.617 (M, G50, outcrossed), as shown in Figure 1. The fixed factors together explained 30.0% of the variance in logit-survival, while the random factors explained 3.6% and the rest (66.3%) was residual variance among individuals (Table 2). Among the random factors block was significant, as well as the replicate*treatment interaction. However, the fraction of variance due to replicate and all its interactions was less than 1%. Among the fixed effects, treatment, line type, and their interaction were significant by LRT (Table 2), meaning that there was significant inbreeding depression, and that line types differed in average juvenile survival, but also in their levels of inbreeding depression. In addition, no consistent change in survival or inbreeding depression was detected between G20 and G50 on average over all line types (experiment effect and experiment*treatment interaction both NS). However, the highly significant line type*experiment interaction revealed strong line type-specific changes in survival.

**Figure 1.**
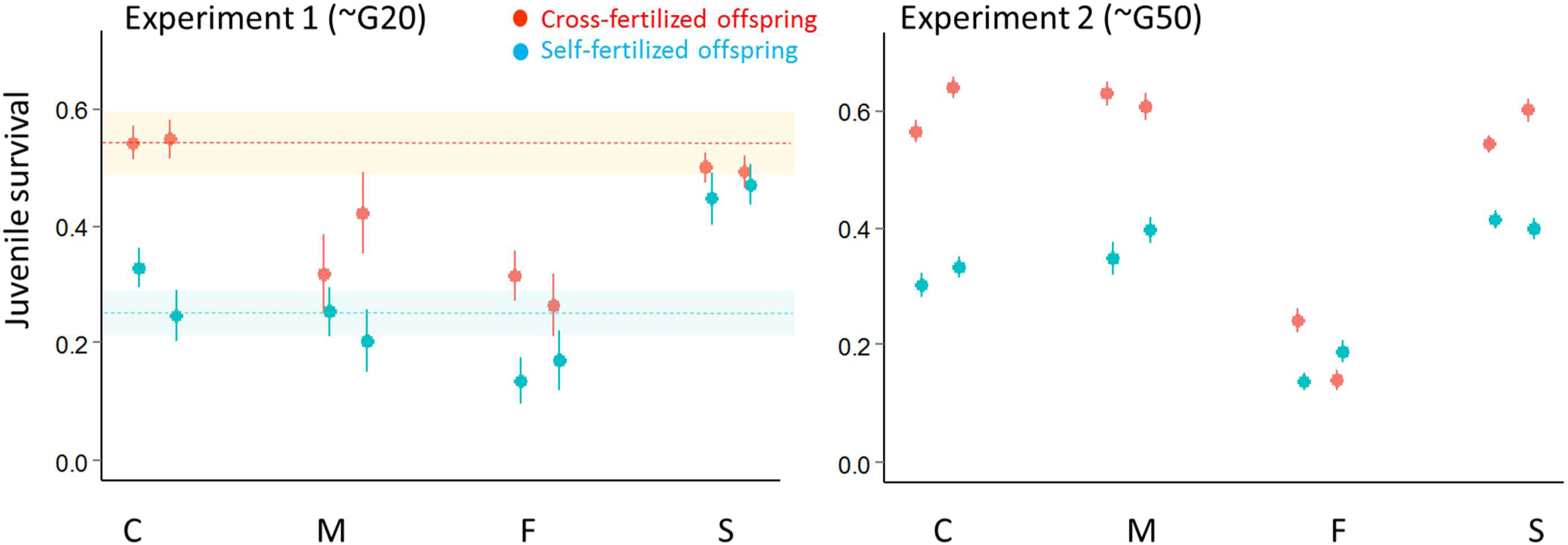
Juvenile survival in the two experiments, for each line and experimental treatment (red: outcrossing, blue: selfing). The average values are given together with their SE over individuals, the two replicates are represented separately for each line type. The relative difference in survival between outcrossed and self-fertilized offspring represents inbreeding depression (δ). The values of δ ±SE for each line type and each experiment are C: 0.46±0.10 (1^st^expt) and 0.47±0.08 (2^nd^expt), F: 0.49±0.15 and 0.16±0.34, M: 0.40±0.13 and 0.40±0.06, and S: 0.08±0.12 and 0.28±0.06. We did not measure juvenile survival in the founding G0 population common to all lines. However we were able to retrieve from a previous study [46] data on the juvenile survival of first-generation self-fertilized and cross-fertilized offspring from 7 of the 10 natural populations that were used to constitute the founding population. The means (±-SE across populations) were respectively : 0.544± 0.054 and 0.251± 0.038 (indicated by dashed lines and shaded areas in the left panel), with an inbreeding depression of δ = 0.524 ± 0.077.

In order to further characterize these effects, we report the average survival of each line type/treatment/experiment, as well as effect sizes and their significance (Wald tests) for each term in the full linear model (Table 3). We chose the C line type, the G20 experiment, and the outcrossed treatment as references in the linear model. The intercept of the model therefore reflected (in logit-scale) the juvenile survival of the C type under outcrossing in G20. The average survival of this reference category was 0.495 (Table 3, Figure 1). Wald tests indicate that C lines exhibited significant inbreeding depression (Treatment (selfed) term) with *ca*. 50% decrease in survival in inbred offspring relative to outbred ones (inbreeding depression estimates are given in the legend of figure 1). The performance of the C lines, irrespective of treatment, did not significantly change between the G20 and G50 experiments (Experiment (G50) and Treatment * Experiment (selfed, G50) in Table 3). In addition the G20 means of C lines were close to those obtained from natural populations that were used to constitute the founding population, measured in a previous study (see Figure 1).

To detail the line type effect and its interaction with treatment and experiment, we will examine the results from each line type (M, F, S) in turn, keeping C as a reference (Table 3 and Figure 1). Outcrossed juveniles from the M line type survived significantly less than those from C in G20 (line type (M) in Table 3), but C and M exhibited similar inbreeding depression (Line type * Treatment, (M,selfed), *NS*). The difference in survival between M and C under outcrossing disappeared in G50 (significant Line type by Experiment interaction (M, G50)), while inbreeding depressions of C and M remained similar (three-way interaction (M, selfed, G50) *NS*).

Among all line types, the F line type presented the most striking differences with controls. They had a much lower juvenile survival than C in the G20 experiment (Line type (F)) and this holded irrespective of treatment, as inbreeding depression did not differ significantly between F and C (Line type * treatment (F,selfed)). The difference in outbred survival rates between F and C became significantly larger in G50 (Line type by Experiment (F, G50), fig. 1) while inbreeding depression remained not significantly different (three-way interaction (F, selfed, G50) *NS*).

Finally, the survival in S lines did not differ significantly from that in C lines under outcrossing in G20 (Line type (S)). However, their performances under inbreeding were higher, as their inbreeding depression was significantly lower (Line type * treatment (S, selfed)). These patterns did not significantly change in time, as the corresponding components of the Line type * experiment (S, G50) and three-way (S, selfed, G50) interactions were *NS*. The figures suggest that inbreeding depression may have increased in the S line type between G20 and G50 (Figure 1) but this change turns out to be *NS*.

Microsatellite genetic diversity varied from 0.14 to 0.53 among the EELs (Figure 2, supplementary Table S1). Within each line type the two replicates were similar, except for the C type (C2 showing reduced variation compared to C1). All M and F EELs had diversities and numbers of alleles comparable to C (*i.e*. within the range between C1 and C2). The S EELs, in contrast, had lower diversity and number of alleles than all other EELs. All *f* values were small and none significantly departed from zero (Table S1)

**Figure 2.**
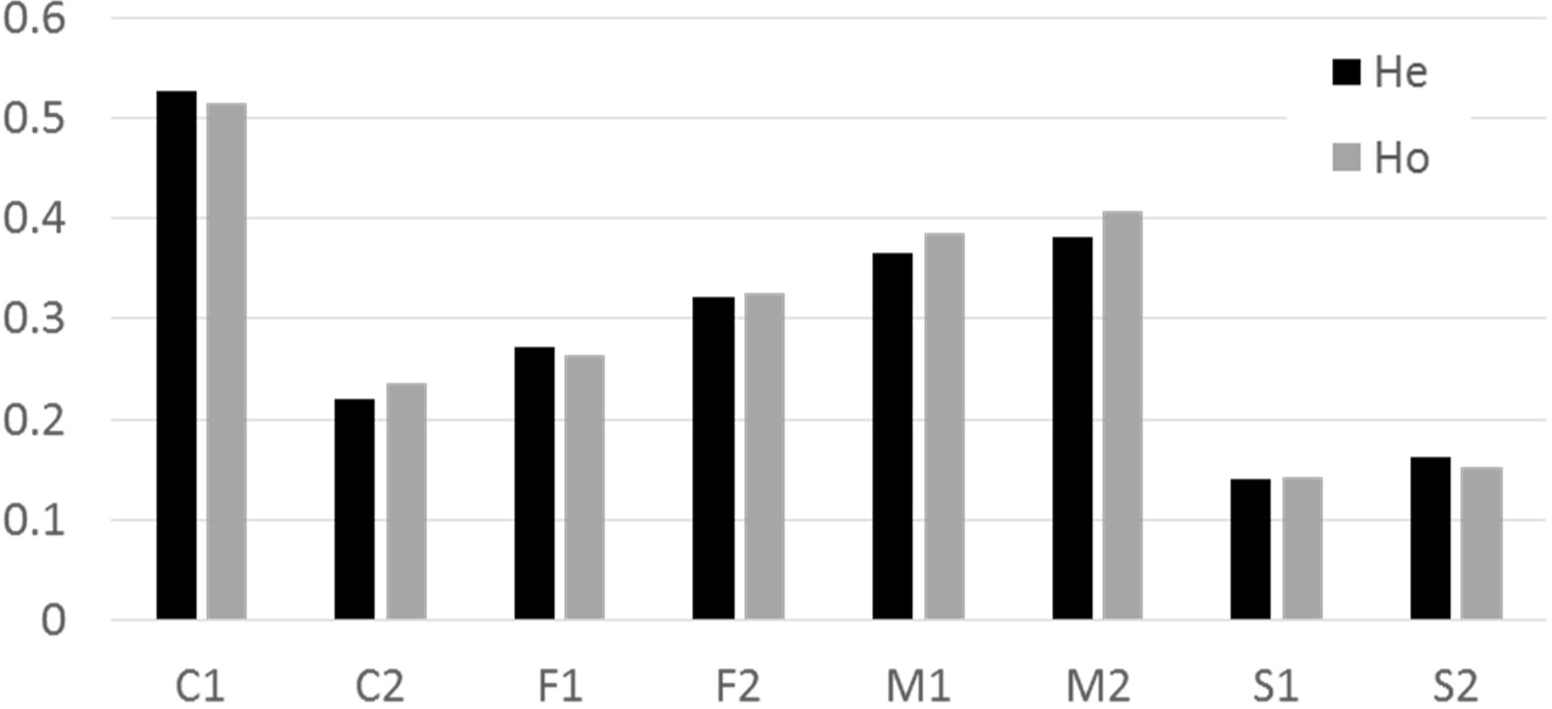
Genetic diversity (expected heterozygosity H_e_) and observed heterozygosity (H_o_) at seven microsatellite loci per line (32 individuals per line) at the 49^th^ generation. Each line type (C, F, M and S) is represented by two replicates.

## Discussion

### Suppression of sexual selection leads to the accumulation of deleterious alleles

In our design, the F lines represent the fate of a population in the absence of sexual selection. This absence had a spectacular effect: juvenile survival dropped to 54% of the controls (C) after 20 generations and down to 32% after 50 generations. The F lines have therefore progressively accumulated deleterious alleles, which may result partly from new mutations, and partly from the increase in frequency of alleles present in the founding population. All our lines were subject to mutation, genetic drift and selection. The controls themselves did not noticeably change in time, with performances in the range of previous studies on wild-caught snails [46], suggesting that they were not far from an equilibrium load. The genetic diversity at (neutral) microsatellite loci was not lower in F than in C lines, suggesting that their effective sizes were not strikingly different – the census sizes were regulated every generation at 90 individuals per line, and the reduced variance in male reproductive success is expected, if anything, to increase the effective size in F lines relative to C, rather than decrease it. The low performances of F lines are therefore likely due to the change in the selection regime on deleterious alleles rather than to an accelerated genetic drift. As predicted by the genic capture hypothesis [6], our results therefore imply that many alleles that are deleterious to juvenile survival in **P. acuta** are normally eliminated by sexual selection through their pleiotropic effects on male success, an effect that is suppressed under monogamy (F lines). In agreement with this idea, a previous study [48] on *Physa acuta*, showed that in conditions allowing male-male competition, inbreeding depression was stronger on male reproductive success than on female reproductive success –suggesting a higher impact of deleterious mutations on male success.

### Effect of sexual selection in simultaneous hermaphroditic <u>vs</u>. gonochoric species

Previous experiments testing the impact of sexual selection through experimental evolution in gonochoric species have yielded inconsistent results, partly reflecting a balance bewteen the opposing effects of sexual selection and sexual conflict on population fitness, that may depend on the study system [14–18]. Our study is the first to address this question in hermaphrodites. Hermaphrodites can differ from gonochoric species in two respects: the intensity of sexual selection and the co-expression of male and female traits in a single individual, which has consequences on sexual conflicts. With regard to the first aspect, several recent studies have shown that strong sexual selection occurs in predominantly outcrossing simultaneous hermaphrodites, for example snails [32,49] and worms [31], contradicting the Darwinian view that they have too low mental powers to undergo sexual selection [50]. In particular, *P. acuta* is an outcrossing and highly polyandrous snail, with intense sexual selection on the male function at both the pre- and post-copulatory stages [30,36]. With regard to sexual conflicts, hermaphrodites differ from gonochoric species because some fitness traits, such as juvenile survival, are necessarily a common component of both male and female fitness. In gonochoric species all fitness traits are expressed separately in either male, or female contexts, and therefore potentially affected by sexual conflicts. In hermaphrodites, the opportunity for such conflicts is high for reproductive traits (e.g., the production of male gametes may be in trade-off with that of female gametes), but low for pre-reproductive survival. Indeed, any reduction in juvenile survival automatically reduces both male and female fitness by the same factor within each individual. The spectacular effect of the suppression of sexual selection in our study may therefore be a consequence of the trait chosen, and the hermaphroditic condition, which reduce the impact of sexual conflicts compared to most studies in gonochoric organisms.

### Sex-specific effects of relaxed selection

In theory, the population benefits of sexual selection are linked to an asymmetry between the sexes. The theory assumes that more deleterious mutations are eliminated through the unsuccessful reproduction of males than through that of females – a cheap way to clear the mutation load, as population growth depends mostly on female fertility [4]. Surprisingly, most experimental evolution studies suppressed sexual selection in males, but did not symmetrically limit the opportunity for selection in females. Some studies, such as [12], limited only the sexual component of selection in females (using male-biased sex ratios). This effectively tests whether sexual selection is stronger in males, but not that variation in male reproductive success is more efficient than variation in female reproductive success at eliminating deleterious mutations. Indeed female reproductive success is determined by fertility and largely independent from the number of mates in many species, including our study system [30]. Thus, limiting the number of males per female is not expected to have a large effect. To validate sexual selection theory, one must therefore show that variation in male fitness (determined by sexual selection) is more efficient at clearing deleterious alleles than variation in female fitness (mostly determined by fertility), and to this end one must relax fertility selection (rather than sexual selection) on females. To our knowledge, a single experimental evolution study conducted in *Drosophila serrata* compared the effects of relaxed sexual selection (monogamy) to those of reduced variation in female reproductive success, using a design similar to ours [15]. Contrary to theoretical expectations, in that study, female selection turned out to be beneficial, and sexual selection costly, to population productivity. This was interpreted as an effect of sexual conflict, linked to the cost of female harassment by males in lines with opportunity for sexual selection [18,51].

Our results, on the contrary, support the theory of purging by sexual selection. Female selection was relaxed in the M lines, and juvenile survival did not decrease on the long term as in F lines, demonstrating that selection on female reproduction was not as important as that on male reproduction to purge the genetic load. Therefore, alleles affecting juvenile survival have stronger pleiotropic effects on male than on female reproduction, as expected from the idea that sexual selection makes male fitness more condition-dependent [6]. We would also expect the performances of M lines to be somewhat reduced compared to controls (although less than in F lines), if some pleiotropic effects on female reproduction existed. A moderate reduction was indeed observed in G20, but it disappeared around the 50^th^ generation, where M lines survived as well as the controls. We did not come up with a clear explanation for this observation. However, our results overall show that the contribution of female reproduction to purging deleterious alleles acting on juvenile survival is much smaller than that of male reproduction. Further studies will be required to examine the evolution of adult reproductive traits in our EELs, especially considering that adult traits might more likely be under the influence of sexual conflict than juvenile survival.

### Inbreeding depression and purging under frequent selfing

In hermaphrodites, complete self-fertilization suppresses, and partial self-fertilization reduces, sexual selection, and the associated genetic purging. However, selfing increases homozygosity and exposes partially recessive alleles to selection, therefore promoting another form of purging [24,25]. This process is central to classical models of mating system evolution, whereby purging in turn selects for higher selfing rates, in a self-reinforcing loop [52]. In line with the hypothesis of purging by inbreeding, inbreeding depression was reduced in S lines, submitted to frequent selfing, compared to C lines in G20, as already pointed out in a previous study [39]. Moreover, M and F lines, which evolved under pure outcrossing like the controls, preserved a large inbreeding depression, not significantly different from that of C. The difference in inbreeding depression between S and all other lines seemed attenuated in G50, but this temporal change is not significant. Overall, inbreeding depression was reduced only in the S lines, and this reduction resulted from an increase in the survival of inbred juveniles, rather than a decrease in the survival of outbred juveniles. Interestingly, microsatellite analysis did not detect hints of self-fertilization among the juveniles produced by nonisolated individuals, even in S lines (all *Fis* were NS). The outcrossed offspring in the S lines were therefore truly outcrossed, and our estimates of inbreeding depression were not downwardly biased (which would have been possible in case of partial selfing). Overall, the results confirm that some purge of deleterious alleles with partially recessive action (i.e. those that contribute to inbreeding depression) has occurred specifically in the lines frequently exposed to selfing.

Two genomic compartments are often considered in the genetic architecture of inbreeding depression: very recessive and strongly deleterious alleles on one hand, and mildly deleterious and weakly recessive alleles on the other hand [25]. The semi-lethal recessives are easily purged under regular selfing, while the slightly deleterious ones are not, making a part of inbreeding depression hard to get rid of [24,53]. In an initially outcrossing population exposed to regular selfing, purging is therefore expected to slow down as the recessive semi-lethals are eliminated and the small-effect alleles persist. Their persistence may also be facilitated by the decrease in the effective size due to both homozygosity and selective interferences under self-fertilization [54], which is confirmed here by the lower genetic diversity of the S lines compared to all others. This persistence might explain that inbreeding depression still persisted at G50.

The classical conception of purging through inbreeding, based on allele recessivity, ignores the potential pleiotropy of deleterious alleles on sexually-selected traits. However, our results in F lines imply that at least some alleles affecting juvenile survival pleiotropically affect male reproduction. The suppression of sexual selection could therefore also relax selection on such alleles in S lines leading to a reduction in the survival of outbred juveniles as in F lines. This was not observed, whether in G20 or in G50. This means that the efficiency of purging through inbreeding is in fact much higher than we could infer based on the comparison between S and C lines (see [39]). The relevant controls are indeed the F rather than the C lines, since S differs from F by self-fertilization only rather than by both self-fertilization and sexual selection (table 1). That S lines maintain high juvenile survival implies that inbreeding does not only eliminate recessive semi-lethals, but also prevents the accumulation of alleles with deleterious effects on both juvenile survival and male reproductive success, that would otherwise take place in the absence of sexual selection. The genic-capture model of sexual selection assumes that the genetic variance in male sexual traits mostly reflects the genomic load, through a condition-dependent reaction norm [6]. Thus, our S lines, if this reaction norm has not itself evolved, may still have the potential to express high male reproductive success if put back in conditions of outcrossing and sexual selection. In consistency with this idea, Noel et al. (2016) [39] did not find any significant reduction in male reproductive success in G20 outbred individuals of the S lines compared to controls. Further studies will be needed to extend to later generations.

Although our data suggest that inbreeding and sexual selection were approximately equally efficient at preserving high juvenile survival, the mutations purged by both processes were not exactly the same. The reduced inbreeding depression in S lines indicates that mutations purged by inbreeding were on average more recessive. Several categories of mutations may contribute to the overall genetic load in our experiment. First, among alleles with recessive deleterious effects on juvenile survival, some may have substantial heterozygous effects on male success (e.g. through adult body condition), and others may lack such effects, thus not being purged by sexual selection. Second, some alleles may not be recessive at all, thus being purged more efficiently by sexual selection. Such alleles would still be sensitive – though only moderately-to inbreeding, as homozygosity increases the additive genetic variance per locus – in our case by a factor (1+*F*_*is*_)=1.5 every second generation [56]. Third, selection against mutations in the haploid phase may contribute to genetic purging, due to competition among male gametes, even during self-fertilization when they come from the same individual (see e.g. [55]). However, as this process occurs under all mating systems including monogamy, it cannot explain differences between selection regimes. All this calls for more studies on the architecture of the genetic load. Some nonlinearities in our data, such as the relative decrease of juvenile survival in M lines in G20, followed by an increase, or the same trend for inbreeding depression in S lines, are also suggestive of a complex genetic architecture of the load, with different compartments purged at different speeds depending on the mechanism.

### Two routes to purging: population consequences

The enhancement of the purging process through sexual selection is one of the forces that may stabilize biparental sexual reproduction against invasion by asexuals [4]. In this study, we have shown that such an advantage can also apply to outcrossing hermaphrodites. This implies that in the latter, self-fertilization may come at a risk of accumulating deleterious alleles, by suppressing sexual selection. Our data however indicate that purging by inbreeding can compensate for the lack of purging by sexual selection, at least at the scale of a few tens of generations, and therefore suggest that the latter does not represent a strong obstacle to an evolutionary transition towards selfing. However, it is not clear whether purging by inbreeding would be efficient at preventing decreases in population fitness over a larger number of generations and with higher selfing rates (*i.e*. nearly obligate selfing instead of the average selfing rate of 50% we imposed). Selective interference and reduced effective size may eventually limit the efficiency of purging (and of selection in general) in predominant selfers [54,57,58], making them evolutionary dead-ends [59,60]. On the other hand, our results suggest the possibility of a “best-of-both-worlds strategy” alternating between episodes of inbreeding (e.g., facultative selfing or mating between relatives) and of outcrossing with strong opportunity for sexual selection, leading to the purging of both the recessive and non-recessive components of the load without the disadvantages of continuous inbreeding. When avoiding mutation accumulation is an issue for yield or conservation, for example in cultivated plants or captive animals, it may be complicated to implement this strategy (e.g., to find the optimal level of inbreeding), but our data at least suggest that random monogamy is probably not a good idea. In natural conditions, hermaphroditic species may experience strong variation in mating or pollination opportunities. In facultative selfers, such as many snails (including *P. acuta*) or plants, this may induce an alternation in mating systems that efficiently keeps deleterious mutations in check.

### Conclusion

Our study provides strong support to the theory of genetic purging by sexual selection. We tested this theory for the first time in hermaphrodites and confirmed that suppressing selection on the female function did not compromise genetic purging as much as suppressing sexual selection on the male function. We also showed that inbreeding was, at least on the short term, a very efficient alternative way to enhance genetic purging. Inbreeding not only eliminates recessive alleles, thus reducing inbreeding depression and preserving the fitness of inbred individuals. It also reduces the fraction of genetic load expressed in outbred individuals, which should otherwise increase because of the absence of sexual selection. This calls for more work on the architecture of the genetic load, and on the optimal strategies to limit its impacts in agronomical and natural contexts.

## Acknowledgements

The authors are grateful to Yohann Chemtob and Tim Janicke for help with the experiments, to the service des marqueurs génétiques at CEFE and the Environmental Genomics platform (GenSeq; LabEx CeMEB) for support with the molecular analyses. This work was performed as part of E. Noël’s PhD project and funded by the French ‘Agence Nationale de la Recherche’ (AFFAIRS ANR-12-ADAP-005, BioAdapt program, and ESHAP ANR-12-BSV7-0015 to P. David), and by CNRS (to P. David and P. Jarne). N. Bonel was supported by CONICET. The authors of this preprint declare that they have no financial conflict of interest with the content of this article.

## Data

Raw data from this study are publicly available at DOI 10.17605/OSF.IO/VNWHB

---

**Figure S1.**
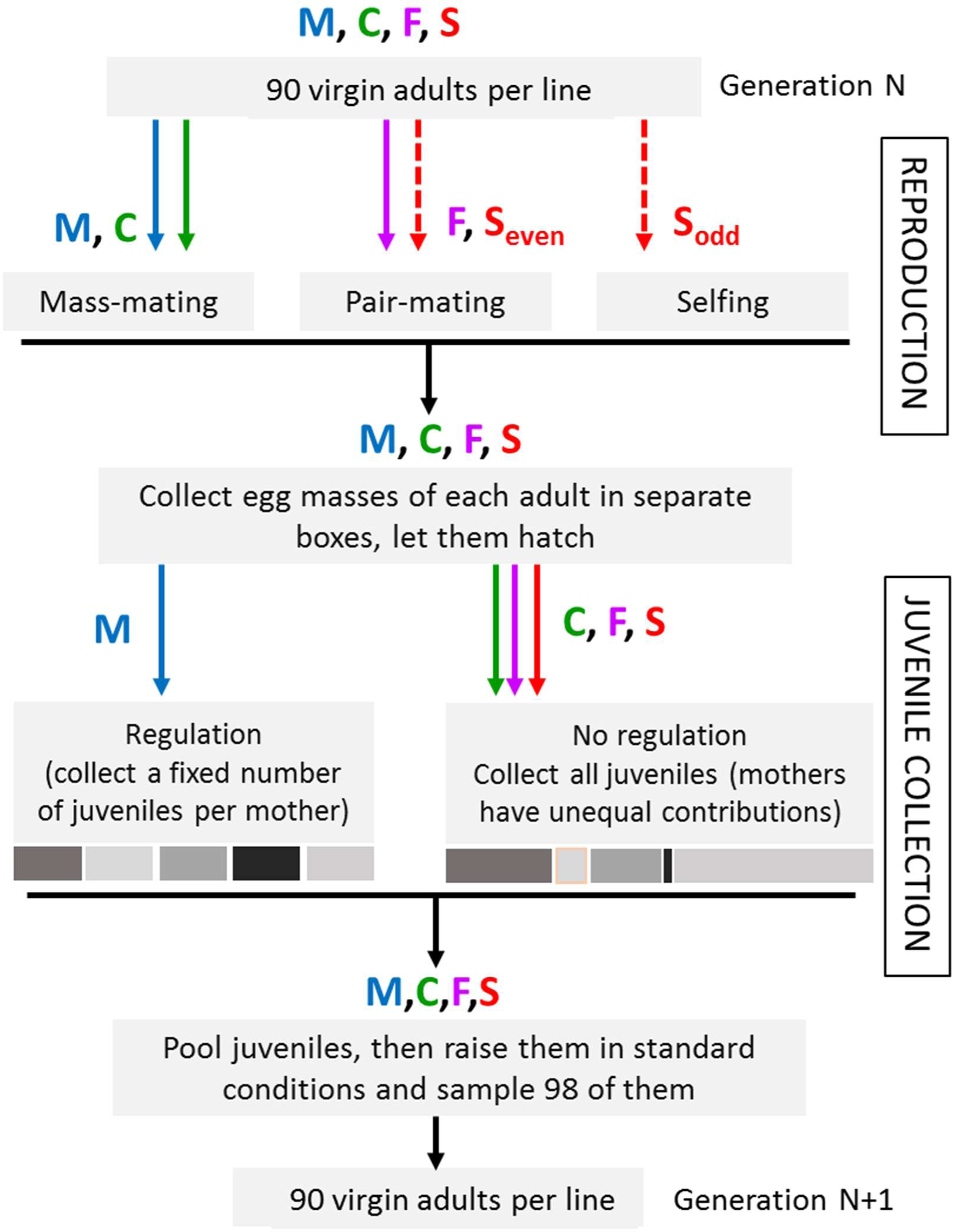
Passage from generation N to generation N+1 in the four types of experimental evolution lines (C; M: F, and S:). All line types have been represented as different paths on the same diagram to stress similarities and differences among them at each stage. Differences arise at the reproduction stage (note that S lines alternate between two modes for even-numbered and odd-numbered generations) and at the juvenile collection stage. The numbers of individuals conserved at each step was calculated in order to obtain a population of approximately 90 reproductive virgin adults at the time of reproduction (this number could vary between 80 and 98 depending on mortality). In practice, when we collected juveniles, we initially kept a large excess of them. In C, F and S lines several hundreds of juveniles were first pooled in a large aquarium then the aquarium was sampled to make 45 boxes of three juveniles (N=135). They then grew for one more week until we sampled 98 of them to raise individually, which ended up at ca. 90 reproductive individuals (80 to 98) given mortality. In M lines, we did not pool juveniles in aquaria and directly made one box of three juveniles for each mother separately (when possible). After growing them one week, we collected one juvenile per box (=i.e. per mother) to raise it in isolation. Because of mortality or failure at the egg-laying or hatching stages, we usually obtained less than 98 maternal broods, so we added a second juvenile taken in some of the boxes at random until we had N=98. This resulted in ca. 90 (80-98) reproductive adults as in the other lines. In all lines, some juveniles in excess were grown in backup aquaria, and served to compensate for mortality if the number of adults turned out to be below 80.

**Table S1.** Microsatellite variation in all experimental evolution lines (N=32 individuals per line, 7 polymorphic loci) at the 49^th^ generation. Each type (C, F, M and S) is represented by two replicate lines. Genetic diversity *H*_*e*_, observed heterozygosity *H*_*o*_, average number of alleles per locus *N*_*all*_, and inbreeding coefficient *f*.

|  | C1 | C2 | F1 | F2 | M1 | M2 | S1 | S2 |
| --- | --- | --- | --- | --- | --- | --- | --- | --- |
| $H_e$ | 0.526 | 0.219 | 0.272 | 0.321 | 0.366 | 0.381 | 0.141 | 0.163 |
| $H_o$ | 0.515 | 0.237 | 0.263 | 0.326 | 0.385 | 0.407 | 0.143 | 0.152 |
| $N_{all}$ | 3.714 | 1.714 | 1.857 | 2.429 | 2.571 | 2.429 | 1.571 | 1.571 |
| $f$ | 0.022 | -0.080 | 0.034 | -0.014 | -0.055 | -0.071 | -0.012 | 0.070 |

## References

1. Eyre-Walker, A., and Keightley, P.D. (2007). The distribution of fitness effects of new mutations. Nat. Rev. Genet. 8, 610–618.

2. Charlesworth, B. (2009). Effective population size and patterns of molecular evolution and variation. Nat. Rev. Genet. 10, 195–205.

3. Agrawal, A.F., and Whitlock, M.C. (2012). Mutation load: the fitness of individuals in populations where deleterious alleles are abundant. Annu. Rev. Ecol. Evol. Syst. 43, 115–135.

4. Agrawal, A.F. (2001). Sexual selection and the maintenance of sexual reproduction. Nature 411, 692–695.

5. Agrawal, A.F. (2011). Are males the more “sensitive”sex? Heredity 107, 20–21.

6. Rowe, L., and Houle, D. (1996). The lek paradox and the capture of genetic variance by condition dependent traits. Proc. R. Soc. London B Biol. Sci. 263, 1415–1421.

7. Hollis, B., Fierst, J.L., and Houle, D. (2009). Sexual selection accelerates the elimination of a deleterious mutant in Drosophila melanogaster. Evolution 63, 324–333.

8. Jarzebowska, M., and Radwan, J. (2010). Sexual selection counteracts extinction of small populations of the bulb mites. Evolution 64, 1283–1289.

9. Clark, S.C.A., Sharp, N.P., Rowe, L., and Agrawal, A.F. (2012). Relative effectiveness of mating success and sperm competition at eliminating deleterious mutations in Drosophila melanogaster. PLoS One 7, e37351.

10. Sharp, N.P., and Agrawal, A.F. (2013). Male-biased fitness effects of spontaneous mutations in Drosophila melanogaster. Evolution 67, 1189–1195.

11. Pélabon, C., Larsen, L.-K., Bolstad, G.H., Viken, A.A., Fleming, I.A., and Rosenqvist, G. (2014). The effects of sexual selection on life-history traits: an experimental study on guppies. J. Evol. Biol. 27, 404–416.

12. Lumley, A.J., Michalczyk, Ł., Kitson, J.J.N., Spurgin, L.G., Morrison, C.A., Godwin, J.L., Dickinson, M.E., Martin, O.Y., Emerson, B.C., Chapman, T., et al. (2015). Sexual selection protects against extinction. Nature 522, 470–473.

13. Singh, A., Agrawal, A.F., and Rundle, H.D. (2017). Environmental complexity and the purging of deleterious alleles. Evolution 71, 2714–2720.

14. Whitlock, M.C., and Agrawal, A.F. (2009). Purging the genome with sexual selection: reducing mutation load through selection on males. Evolution 63, 569–582.

15. Rundle, H.D., Chenoweth, S.F., and Blows, M.W. (2006). The roles of natural and sexual selection during adaptation to a novel environment. Evolution 60, 2218–2225.

16. Michalczyk, Ł., Millard, A.L., Martin, O.Y., Lumley, A.J., Emerson, B.C., and Gage, M.J.G. (2011). Experimental evolution exposes female and male responses to sexual selection and conflict in Tribolium castaneum. Evolution 65, 713–724.

17. Arbuthnott, D., and Rundle, H.D. (2012). Sexual selection is ineffectual or inhibits the purging of deleterious mutations in Drosophila melanogaster. Evolution 66, 2127–2137.

18. Chenoweth, S.F., Appleton, N.C., Allen, S.L., and Rundle, H.D. (2015). Genomic evidence that sexual selection impedes adaptation to a novel environment. Curr. Biol. 25, 1860–1866.

19. Radwan, J. (2004). Effectiveness of sexual selection in removing mutations induced with ionizing radiation. Ecol. Lett. 7, 1149–1154.

20. McGuigan, K., Petfield, D., and Blows, M.W. (2011). Reducing mutation load through sexual selection on males. Evolution 65, 2816–2829.

21. Almbro, M., and Simmons, L.W. (2014). Sexual selection can remove an experimentally induced mutation load. Evolution 68, 295–300.

22. Grieshop, K., Stångberg, J., Martinossi-Allibert, I., Arnqvist, G., and Berger, D. (2016). Strong sexual selection in males against a mutation load that reduces offspring production in seed beetles. J. Evol. Biol. 29, 1201–1210.

23. Byers, D.L., and Waller, D.M. (1999). Do Plant Populations Purge Their Genetic Load? Effects of Population Size and Mating History on Inbreeding Depression. Annu. Rev. Ecol. Syst. 30, 479–513.

24. Crnokrak, P., and Barrett, S.C.H. (2002). Perspective: purging the genetic load: a review of the experimental evidence. Evolution 56, 2347–2358.

25. Charlesworth, D., and Willis, J.H. (2009). The genetics of inbreeding depression. Nat. Rev. Genet. 10, 783–796.

26. Hartfield, M., Bataillon, T., and Glémin, S. (2017). The Evolutionary Interplay between Adaptation and Self-Fertilization. Trends Genet. 33, 420–431.

27. Leonard, J.L. (2006). Sexual selection: lessons from hermaphrodite mating systems. Integr. Comp. Biol. 46, 349–367.

28. Delph, L.F., and Ashman, T.-L. (2006). Trait selection in flowering plants: how does sexual selection contribute? Integr. Comp. Biol. 46, 465–472.

29. Moore, J.C., and Pannell, J.R. (2011). Sexual selection in plants. Curr. Biol. 21, R176–R182.

30. Pélissié, B., Jarne, P., and David, P. (2012). Sexual selection without sexual dimorphism: Bateman gradients in a simultaneous hermaphrodite. Evolution 66, 66–81.

31. Marie-Orleach, L., Janicke, T., Vizoso, D.B., David, P., and Schärer, L. (2016). Quantifying episodes of sexual selection: Insights from a transparent worm with fluorescent sperm. Evolution 70, 314–328.

32. Hoffer, J.N.A., Mariën, J., Ellers, J., and Koene, J.M. (2017). Sexual selection gradients change over time in a simultaneous hermaphrodite. Elife 6, e25139.

33. Baker, H.G. (1955). Self-compatibility and establishment after’long-distance’dispersal. Evolution 9, 347–349.

34. Escobar, J.S., Cenci, A., Bolognini, J., Haudry, A., Laurent, S., David, J., and Glémin, S. (2010). An integrative test of the dead-end hypothesis of selfing evolution in Triticeae (Poaceae). Evolution 64, 2855–2872.

35. Escobar, J.S., Auld, J.R., Correa, A.C., Alonso, J.M., Bony, Y.K., Coutellec, M.-A., Koene, J.M., Pointier, J.-P., Jarne, P., and David, P. (2011). Patterns of mating-system evolution in hermaphroditic animals: Correlations among selfing rate, inbreeding depression, and the timing of reproduction. Evolution 65, 1233–1253.

36. Pélissié, B., Jarne, P., Sarda, V., and David, P. (2014). Disentangling precopulatory and postcopulatory sexual selection in polyandrous species. Evolution 68, 1320–1331.

37. Janicke, T., David, P., and Chapuis, E. (2015). Environment-dependent sexual selection: Bateman’s parameters under varying levels of food availability. Am. Nat. 185, 756–768.

38. Tsitrone, A., Jarne, P., and David, P. (2003). Delayed selfing and resource reallocations in relation to mate availability in the freshwater snail Physa acuta. Am. Nat. 162, 474–488.

39. Noël, E., Chemtob, Y., Janicke, T., Sarda, V., Pélissié, B., Jarne, P., and David, P. (2016). Reduced mate availability leads to evolution of self-fertilization and purging of inbreeding depression in a hermaphrodite. Evolution 70, 625–640.

40. Jarne, P., Perdieu, M., Pernot, A., and Delay, B. (2000). The influence of self-fertilization and grouping on fitness attributes in the freshwater snail Physa acuta: population and individual inbreeding depression. J. Evol.

41. Tsitrone, A., Duperron, S., and David, P. (2003). Delayed selfing as an optimal mating strategy in preferentially outcrossing species: theoretical analysis of the optimal age at first reproduction in relation to mate availability. Am. Nat. 162, 318–31.

42. Bates, D., Mächler, M., Bolker, B., and Walker, S. (2014). Fitting linear mixed-effects models using lme4. arXiv Prepr. arXiv1406.5823.

43. Elston, D.A., Moss, R., Boulinier, T., Arrowsmith, C., and Lambin, X. (2001). Analysis of aggregation, a worked example: numbers of ticks on red grouse chicks. Parasitology 122, 563–569.

44. Benjamini, Y., and Hochberg, Y. (1995). Controlling the false discovery rate: a practical and powerful approach to multiple testing. J. R. Stat. Soc. Ser. B, 289–300.

45. Sourrouille, P., Debain, C., and Jarne, P. (2003). Microsatellite variation in the freshwater snail Physa acuta. Mol. Ecol. Notes 3, 21–23.

46. Escobar, J.S., Nicot, A., and David, P. (2008). The different sources of variation in inbreeding depression, heterosis and outbreeding depression in a metapopulation of Physa acuta. Genetics 180, 1593–1608.

47. Belkhir, K., Borsa, P., Chikhi, L., Raufaste, N., and Bonhomme, F. (2004). Genetix 4.05. Population Genetics Software for Windows TM, University of Montpellier.

48. Janicke, T., Vellnow, N., Sarda, V., and David, P. (2013). Sex-Specific Inbreeding Depression Depends On The Strength Of Male-Male Competition. Evolution 67, 2861–2875.

49. Anthes, N., David, P., Auld, J.R., Hoffer, J.N.A., Jarne, P., Koene, J.M., Kokko, H., Lorenzi, M.C., Pélissié, B., Sprenger, D., et al. (2010). Bateman gradients in hermaphrodites: An extended approach to quantify Sexual selection. Am. Nat. 176, 249–263.

50. Schärer, L. (2009). Tests of sex allocation theory in simultaneously hermaphroditic animals. Evolution 63, 1377–1405.

51. Rundle, H.D., Chenoweth, S.F., and Blows, M.W. (2009). The diversification of mate preferences by natural and sexual selection. J. Evol. Biol. 22, 1608–1615.

52. Lande, R., and Schemske, D.W. (1985). The evolution of self-fertilization and inbreeding depression in plants. I. Genetic models. Evolution 39, 24–40.

53. Porcher, E., and Lande, R. (2005). The evolution of self-fertilization and inbreeding depression under pollen discounting and pollen limitation. J. Evol. Biol. 18, 497–508.

54. Hartfield, M., and Glemin, S. (2016). Limits to adaptation in partially selfing species. Genetics 203, 959–974.

55. Tazzyman, S.J., Seymour, R.M., and Pomiankowski, A. (2012). Fixed and dilutable benefits: female choice for good genes or fertility. Proc. R. Soc. B 279, 334–340.

56. Falconer, D.S., and Mackay, T.C. (1989). Introduction to quantitative genetics. (John Willey and Sons Inc., New York).

57. Lande, R., and Porcher, E. (2015). Maintenance of quantitative genetic variance under partial self-fertilization, with implications for evolution of selfing. Genetics 200, 891–906.

58. Noël, E., Jarne, P., Glémin, S., MacKenzie, A., Segard, A., Sarda, V., and David, P. (2017). Experimental Evidence for the Negative Effects of Self-Fertilization on the Adaptive Potential of Populations. Curr. Biol. 27, 237–242.

59. Igic, B., and Busch, J.W. (2013). Is self-fertilization an evolutionary dead end? New Phytol. 198, 386–397.

60. Goldberg, E.E., Kohn, J.R., Lande, R., Robertson, K.A., Smith, S.A., and Igić, B. (2010). Species selection maintains self-incompatibility. Science 330, 493–495.

